# Direct Electrical Detection of sub-aM DNA Concentrations

**DOI:** 10.1101/2020.04.03.023176

**Authors:** Maoxiang Guo, Narayanan Madaboosi, Felix Neumann, Mats Nilsson, Wouter van der Wijngaart

## Abstract

Sensors for detecting ultra-low concentrations are limited by the efficient transport of target molecules from large sample volumes to small detection regions. We here report a small-format digital DNA sensor in the shape of a microporous membrane that electrically detects DNA substrates with a concentration as low as 790 zM. This ultra-high sensitivity follows from optimising the mass transport of target DNA to specific receptors on the membrane across multiple spatial scales. mm-sized membranes support the rapid convection of a large sample volume to the detection zone; µm-sized pores ensure that DNA diffusion to the surface-based receptors dominates over convective loss through the pores (low Péclet number), and; at the nm-scale, target-receptor binding dominates over diffusive transport (high Damköhler number). After their efficient capture, the DNA molecules are converted with high specificity into trans-membrane gold nanowires that are detected using a simple, high signal-to-noise, electrical resistance measurement. This sensor design is of interest for detecting low-abundant target molecules without the need for sample amplification or up-concentration, and the mass-transport strategy could be adapted to other surface-based sensing schemes.

---

Detecting low-abundant DNA is of interest in life sciences and biomedicine, for example to early diagnose disease and to detect rare mutations. Single molecule detection (SMD) is the “Holy Grail” of analytical chemistry, as it is the highest resolution measurement one can make. However, the critical measurement and the biggest challenge is the concentration: to detect single molecules in a low concentration solution demands to interrogate large volumes of solution (1 aM ≈ 6 molecules per 10 µL).^1^

Unfortunately, sensitive detectors typically feature small detection zones. Free field optical detectors limit the interrogation volume to reduce background, for example by illuminating only limited sections of the sample using a laser sheet in fluorescence-based detection^2^ or by observing only the evanescent field region in total internal reflection (TIR)-based methods.^3,4^ Acoustic (quartz crystal microbalance, ^5^ surface acoustic wave-based^6^), electrochemical^7^ or photonic (surface plasmon, ^8^ photonic waveguide-based^9^) molecular sensors rely on the interaction of particles with the detector surface, which defines an extremely confined detection volume. The continuous interrogation of small volumes can address the mismatch between the volume of the sample and that of the detection zone in flow-based SMD systems, but this is impractical for low concentrations due to the excessive measurement time, with high resolution fluorescent molecule detection in microchannels reported at flow rates of 0.45 nL·s^−1^.^10^ Another strategy is to actively transport the target molecules through the sample to the detection zone where they create locally an elevated concentration. Electrokinetic transport and trapping mechanism, such as dielectrophoretic, ^11^ isotachophoretic^12,13^ and AC electro-osmotic^14^ techniques, are relatively easy to implement, but cannot be readily combined with SMD. Magnetic bead-based transport allows for the detection of sub-pM protein samples on TIR-based detectors, ^4^ and of 350 zM samples using digital ELISA in fL well arrays.^15^

Several schemes have been reported for the direct electrical detection of DNA, for example, by surface charge perturbation,^17^ by conductance change in semiconductor nanowires,^18^ or by converting DNA to metallic nanowires by stretching the strands between the tips of nanotweezers in solution,^16^ on flat surfaces,^19^ or across porous membranes^20^ followed by decorating them with metallic nanoparticles.

In this work, we explore the highly-specific direct electrical detection of ultra-low concentrations of DNA using a porous membrane-based detector. The low limit of detection (LoD) results from the detector geometry allowing efficient transport of target molecules from the sample bulk to the detection zone. The optimized transport is combined with an efficient and highly-specific DNA-to-gold nanowire conversion, and a high signal-to-noise electrical resistance measurement.

## Results

We built 25 × 75 mm^2^-sized detectors that contain three sample areas with *N* = 6 detection wells each (Fig. 1). Each well is divided in half by a 10 µm thick horizontal porous membrane with 1 µm diameter vertical pores. The top and bottom surfaces of the membrane are coated with gold and functionalized with target-specific single-stranded DNA capture probes.

**Figure 1:**
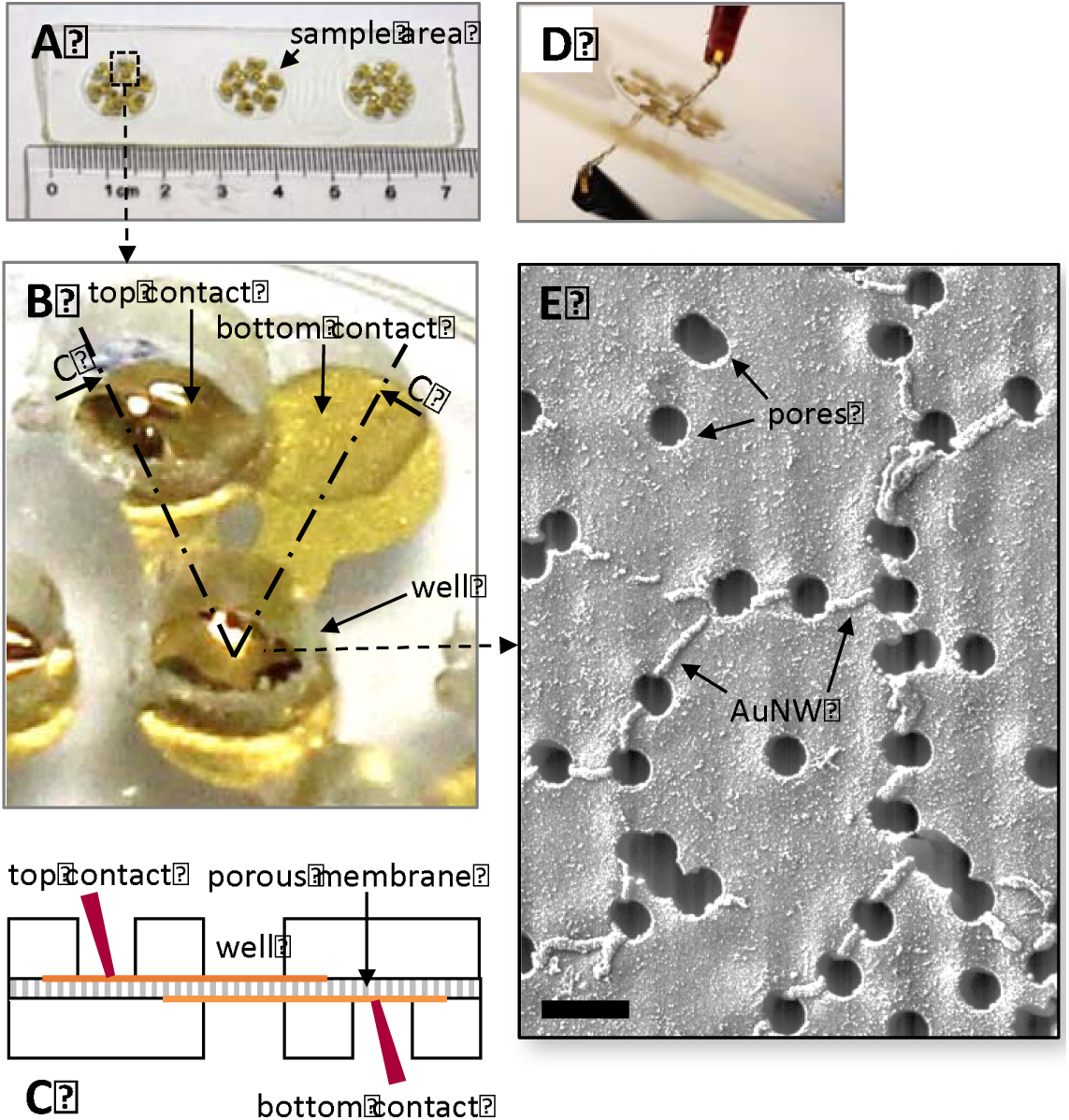
The detector. A) Photograph of a detector with three sample areas that each contain N = 6 measurement wells. B) Close-up photograph and C) schematic cross-section of one measurement well and its top and bottom contact area. D) Photograph of the measurement of the trans-membrane electrical resistance of one well. E) SEM image of the top surface of the membrane after processing of 790 fM DNA sample (scale bar 2 µm).

The assay is illustrated in Fig. 2. During operation, 50 μL of sample is added to one sample area, and positive (50 μL of 790 aM DNA) and negative (50 μL of ≤ 79 zM DNA) control samples to the other two areas. The target DNA in the sample are padlock probes, specifically designed ssDNA oligonucleotides that bind to their capture probes in a circular fashion.^21^ The sample flows through the membrane, driven by gravity, during which process target DNA binds to the surface-based probes. Thereafter, a target-specific RCA assay elongates the captured strands in ssDNA concatemers. A highly specific ligation, which offers single nucleotide discrimination, circularizes the targets. The circularized targets form primers for the subsequent isothermal amplification reaction that converts them into long ssDNA concatemers.^22^ The concatemers are subsequently stretched through the pores and decorated with specifically modified gold nanoparticles, whereafter a gold enhancement connects the nanoparticles to form trans-membrane gold nanowires. SEM imaging confirmed the presence of gold nanowires on the top surface of the membranes. Thereafter, the electrical resistance across the membranes is measured for every well (Table S3). Wells in which gold nanowires result in a finite detectable resistance are defined positive; wells with infinite resistance (open circuit) are defined negative.

**Figure 2:**
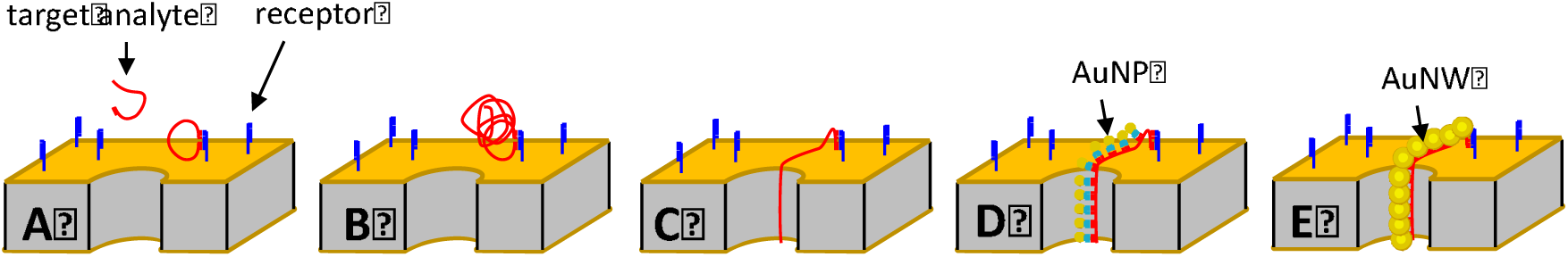
**Schematic of the detector operation**, shown on a section of the polycarbonate membrane with one pore, coated with Ti/Au and functionalized with specific DNA receptors, during sample incubation. A) sample incubation. B) RCA forms elongated DNA concatemers. C) A receding liquid–air interface stretches the concatemers through the membrane pores. D) Gold nanoparticles functionalised with specific DNA oligonucleotides hybridize with the concatemers. E) Gold enhancement of the AuNPs attached to the stretched concatemers form trans-membrane gold nanowires that electrically connect the top and bottom surface of the membrane.

The detector response versus the concentration of active substrate is plotted in Fig. 3. The measurement results underline the ultra-high sensitivity (790 zM detected) and the digital response of the developed detector. The detector provides both ultra-high sensitive and highly specific measurements. To demonstrate the sensor specificity, we chose padlock probes in combination with RCA. ^22^ The assay specificity was confirmed using relevant controls in the assay design (Table S4). Mismatching DNA target or mismatching receptors both resulted in open circuit responses. Removing the DNA receptor or omitting AuNPs from the assay similarly resulted in open circuit responses. These control measurements further confirm the high specificity of our detector by maintaining the required assay stringency.

**Figure 3.**
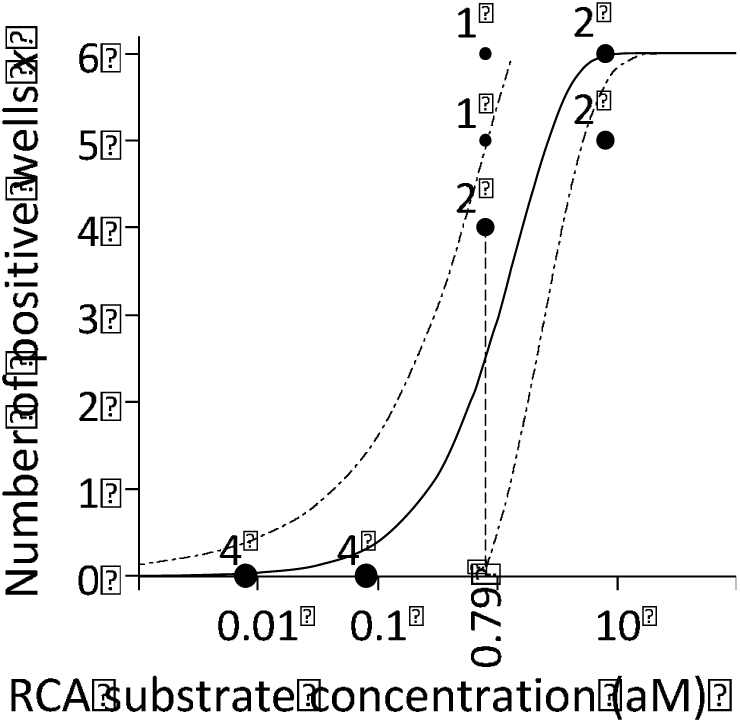
Measurements of RCA substrate (dots) and the theoretical detector response curve 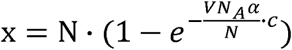 (solid line) for the best-fit value α = 13.7 %. The measurement dot size and numbers beside them indicate the number of times this result was measured. The dash-dotted lines indicate the 2σ interval around the response curve. The dashed line indicates the lowest concentration measured, 0.79 aM.

## Discussion

The main challenge in developing ultra-sensitive detectors remains the sample transport to the detector interrogation zone. Our detector is designed for rapid active sample transport to the interrogation zone with low loss of analyte by convection, and has therefore explicit advantages for the timely detection of ultra-low concentrations. Below, we discuss the analyte transport, electrical resistance values, the detector performance, and the application potential of the detector. Table 1 lists the variables and the governing parameters and their measured or estimated values.

**Table 1:** Variables and parameters and their (estimated) values.

| Parameter / Variable | Defntion | Experimental value |
| --- | --- | --- |
| $\alpha$ | the probability of converting a DNA target molecule in the sample into a detectable gold nanowire | $\approx 13.7\%$ |
| $\eta$ | the sample viscosity | $\approx 10^{-3} \text{ Pa.s}$ |
| $\lambda$ | the expected number of detectable gold nanowires per well $\frac{m\alpha}{N}$ | |
| $A$ | the active detector membrane area of one well | $2 \text{ mm}^2$ |
| $c$ | the sample concentration | |
| $c_s$ | the surface concentration of the receptors | $> 10^{-8} \text{ mol/m}^2$ <sup>(23,24)</sup> |
| $D$ | the diffusion coefficient of the analyte in the sample | $\approx 4.3 \cdot 10^{-11} \text{ m}^2/\text{s}$ <sup>(25)</sup> |
| $Da$ | the Damköhler number for analyte transport at the membrane pore entrance $\frac{k_{on} \cdot c_s \cdot R_p}{D}$ | $> 10^3$ |
| $f$ | the accepted failure rate when defining the detector LoD or saturation level | |
| $k_{on}$ | the binding reaction rate of the analyte to the surface bound receptors | $10^6 \dots 10^7 \text{ M}^{-1}\text{s}^{-1}$ <sup>(26,27)</sup> |
| $L_p$ | the pore length | $10 \text{ }\mu\text{m}$ |
| $m$ | the copy number of target analyte molecules $c \cdot V \cdot N_A$ | |
| $N$ | the number of detector wells | 6 |
| $N_A$ | the Avogadro number | $\approx 6.022 \cdot 10^{23}$ |
| $P$ | the pressure drop over the membrane | $\approx 3 \text{ mm H}_2\text{O}$ |
| $Pe$ | the Péclet number for analyte transport at the membrane pore entrance $\frac{u_p \cdot R_p}{D}$ | $\approx 1$ |
| $R_p$ | the pore radius | $0.5 \text{ }\mu\text{m}$ |
| $T$ | the absolute temperature at which the detection takes place | $\approx 297 \text{ K}$ |
| $t_{conv}$ | the time for converting trapped DNA to gold nanowires | 170 min |
| $t_{trans}$ | the time for transporting DNA from the sample bulk to the membrane surface | 35 min |
| $u_p$ | the average convective sample velocity at the pore entrance | |
| $V$ | the sample volume | $50 \text{ }\mu\text{L}$ |
| $x$ | the number of positive detector wells | |

The transport of target molecules from their dissolved state in the bulk of the sample to a bound state on the top surface of the membrane involves three subsequent transport modes: rapid convection of the sample towards (and through) the membrane; diffusion of analyte to the top surface of the membrane prior to entering the membrane pores, and; binding of the analyte to the surface.^28^ Efficient analyte molecule transport requires that, in the vicinity of the pore entrance, the rate of diffusive transport of molecules to the membrane surface is larger than that of the convective transport through the pores, i.e., the Péclet number 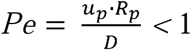, and that the rate of analyte binding to the surface is larger than that of the diffusive transport, i.e., the Damköhler number 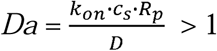. We can always fulfill *Da* > 1 by choosing a large enough value of *R*_*p*_. Using the Hagen-Poiseuille equation 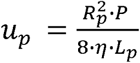 for laminar flow through the pores, we can rewrite 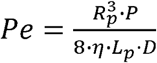. We can always fulfill *Pe* < 1 by choosing the trans-membrane pressure drop P sufficiently low, the membrane thickness *L*_*p*_ sufficiently high, or the temperature *T* sufficiently high (Einstein’s theory of Brownian motion^29^ predicts *D* · η ∼ *T* ^−1^). In our experiments, *Pe* ≈ 1, which is sufficient, but not optimal, as a non-negligible fraction of analyte is expected to flow through the pores rather than bind to the membrane.

The measured electrical resistance values have a large variation between the wells (Table S3). We ascribe these resistance variations to local variations in flow velocity between wells and between pores during gold enhancement, leading to variations in the gold deposition rate. The resistance measurements can therefore not resolve the number of trans-membrane wires, and we, therefore, interpret the resistance readout as a digital measure, with the wells being either positive (finite resistance, ≥ 1 wire) or negative (open circuit, 0 wires). The electrical resistance measurements provide an exceptionally high S/N: all experiments with zero sample concentration resulted in zero false-positives (n > 36). Hence, there is no need for defining a readout signal threshold, as in fluorescence detection methods.

A sample area has *N* = 6 individually readable wells and is thus digital. We assume that the successful conversion of a DNA molecule into an electrically detectable gold nanowire is a stochastic process with probability α. For a given molecule copy number *m* in the sample, the number of gold nanowires per well follows a Poisson distribution with an expected number of detectable wires per well A = m · α /N. The response curve, detection limit, and saturation of such sensors are described in detail elsewhere.^30^ The probability distribution for the number of positive wells follows a binomial distribution with mean *N* (1 − *e*^−λ^) and variance σ^2^ = *N* (*e*^−λ^ −*e*^−2λ^), and the best estimation of *m* may, therefore, be calculated from the number of positive wells *x* as 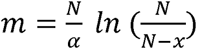. Best parameter fitting allows estimating α≈ 13.7% (Fig.1). We speculate that losses (α < 1) result from non-optimal sample transport (*Pe* ≮ 1), imperfect stretching of concatemers and breaking of DNA concatemers, or an insufficient amount of gold nanoparticles binding to the concatemers. The minimum detectable molecule copy number, *m*_*LoD*_, and the maximum number of molecules quantifiable, *m*_*sat*_, depend on the choice of an acceptable failure rate, *f*, and can be estimated as *m*_*LoD*_ = -ln (f)/α and 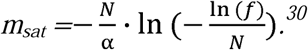 For example, at *f* = 10 %, *m*_*LoD*_ ≈ 17 (limit of detection (LoD) ∼570 zM) and *m*_*sat*_ ≈ 41; at *f* = 50 %, *m*_*LoD*_ ≈ 5 (LoD ∼ 170 zM) and *m*_*sat*_ ≈ 93. The lowest concentration measured with the detector, 790 zM, is almost four magnitude orders below those of previously demonstrated direct electrical DNA measurements. ^20^ We attribute this to the tuned mass transport and the use of dimethyl sulfoxide (DMSO) buffer, instead of phosphate-buffered saline (PBS), during the stretching of the DNA strands, where DMSO is known to decrease self-complementarity reactions of the ssDNA strands, ^31^ thus increasing the likelihood of successful stretching. The dynamic range (∼ *m*_*sat*_*/m*_*LoD*_) is low for the realized detectors but can be increased readily by increasing *N*. The detector has a very simple geometry, and the electrical resistance measurement allows for simple signal acquisition. We thus speculate that the detector can be readily implemented in a silicon MEMS-based format with integrated readout electronics in conjunction with every membrane section. Such implementation would allow a large number of miniaturised wells and a resulting dramatic increase of the dynamic range.

We here demonstrate that a sample area with active membrane area N · A = 12 mm^2^ can detect 790 zM DNA in a 50 μL sample within a detection time t_trans_ + t_conv_ = 35 min +170 min. 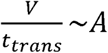 · *u*_*p*_ expresses the trade-off between the detector size (∼A), sample volume, *V*, and the measurement time (*Pe* < 1 limits *u*_*p*_). Optimising the individual assay steps may further reduce the DNA-to-gold nanowire conversion time, *t*_*conv*_.

The overall detector specificity is excellent because it builds on three independent specific events: the target-receptor binding, the ligase binding (the ligase has a specific footprint at both sides of the ligation site that cannot tolerate mismatches), and the AuNP hybridization. The high specificity was confirmed on different assay levels as presented.

RCA allows for a very high level of multiplexing,^32–34^ and therefore, as suggested elsewhere,^20^ we believe that the detector format is suitable for adaptation to multiplexed assays by selectively activating specific wells with specific receptors. Moreover, different assay formats that include target circularization, as in the case of molecular inversion probes, ^32^ offers the required tunability for the platform without compromising the specificity and sensitivity features of RCA. For a given measurement time and LoD, such schemes would require linearly scaling the sample volume and active membrane area with the level of multiplexing, similarly as in multiplexing schemes for digital ELISA.^35^ Furthermore, the membrane-structured detector geometry lends itself for used in conjunction with other transduction principles such as evanescent field photonic sensors.

We here demonstrate the detection of DNA at ultra-low concentrations with high specificity. As suggested elsewhere,^20^ the detector format can also be suitable for adaptation to a combinatorial protein read-out system by adopting the proximity ligation assay.^36–39^ We believe that the performance characteristics, specifically the small size and the simplicity of the design, make the detection principle promising for applications at the point-of-care. Functionalization with relevant pathogen/biomarker-specific probes could make such detectors suitable for application as a powerful diagnostic tool in clinical settings, for example, for the detection of different pathogens,^37^ antibiotic resistance markers,^38^ cancer mutations, and *in situ* transcriptome analysis.^39^

## Conclusion

In conclusion, this work demonstrates how a carefully designed mass transport allows the timely direct electrical detection of ultra-low concentrations of DNA. Compared to state-of-the-art ultra-sensitive detectors, our detector features a competitive LoD and detection time; allows for adaptation to a high dynamic range and level of multiplexing, and; combines a high specificity and signal-to-noise ratio of the readout signal with a simple construction and operation. We expect that the combination of these features can help to bring sensitive digital bioassay formats to use in point-of-care applications.

## Acknowledgement

MG acknowledges the financial support from the China Scholarship Council in China. FN, NM and MN acknowledge the European Union’s Horizon 2020 research and innovation program New Diagnostics for Infectious Diseases (ND4ID) (grant number 675412) for their financial support. We also express our gratitude to Cecilia Aronsson and Mikael Bergqvist for the help with the sample preparation and set-up fabrication, and to Iván Hernández-Neuta for his aid in the lab.

## Materials and methods

### Materials

Porous polycarbonate membranes with a diameter of 13 mm, a thickness of 10 um and containing straight pores of diameter 1 um and pore surface density of 1.4 · 10^7^ cm^-2^ were purchased from Whatman, Florham Park, Morris County, NJ, USA. Off-Stoichiometry thiol-ene-epoxy (OSTE+) was obtained from Mercene Labs AB, Stockholm, Sweden. PDMS was purchased from The Dow Chemical Company, MI, USA. Poly(methyl methacrylate) (PMMA) was purchased from Nordbergs Tekniska AB, Vallentuna, Sweden. T4 DNA ligase, ATP, and dNTPs were purchased from Blirt S.A., Gdansk, Poland. Phi29 polymerase was purchased from Monserate Biotechnology Group, San Diego, CA, USA. Phi29 polymerase buffer was purchased from Thermo Scientific, Waltham, MA, USA. The NAP-5 column, Dimethyl sulfoxide (DMSO), Dithiothreitol (DTT), thiolated poly(ethylene glycol) (PEG), sodium dodecyl sulfate (SDS) were purchased from Sigma-Aldrich, St. Louis, MO, USA. Tris-HCl, PBS, and NaCl were obtained from Karolinska Institute Substrat, Stockholm, Sweden. Bovine serum albumine (BSA) was provided by New England Biolabs, Ipswich, MA, USA. Streptavidin gold nanoparticle solution containing 10 nm diameter particles was purchased from BBI Solutions Company, Crumlin, UK. GoldEnhance solution was purchased from Nanoprobes Inc., Yaphank, NY, USA. Table 2 lists the oligonucleotide sequences of the target hybridization sites of the capture oligonucleotides, the padlock probes, and the AuNPs tag sequences. Milli-Q water was prepared in-house with a resistance > 18 MK. All oligonucleotide were purchased from Integrated DNA Technologies, San Diego, CA, USA.

**Table 2:**
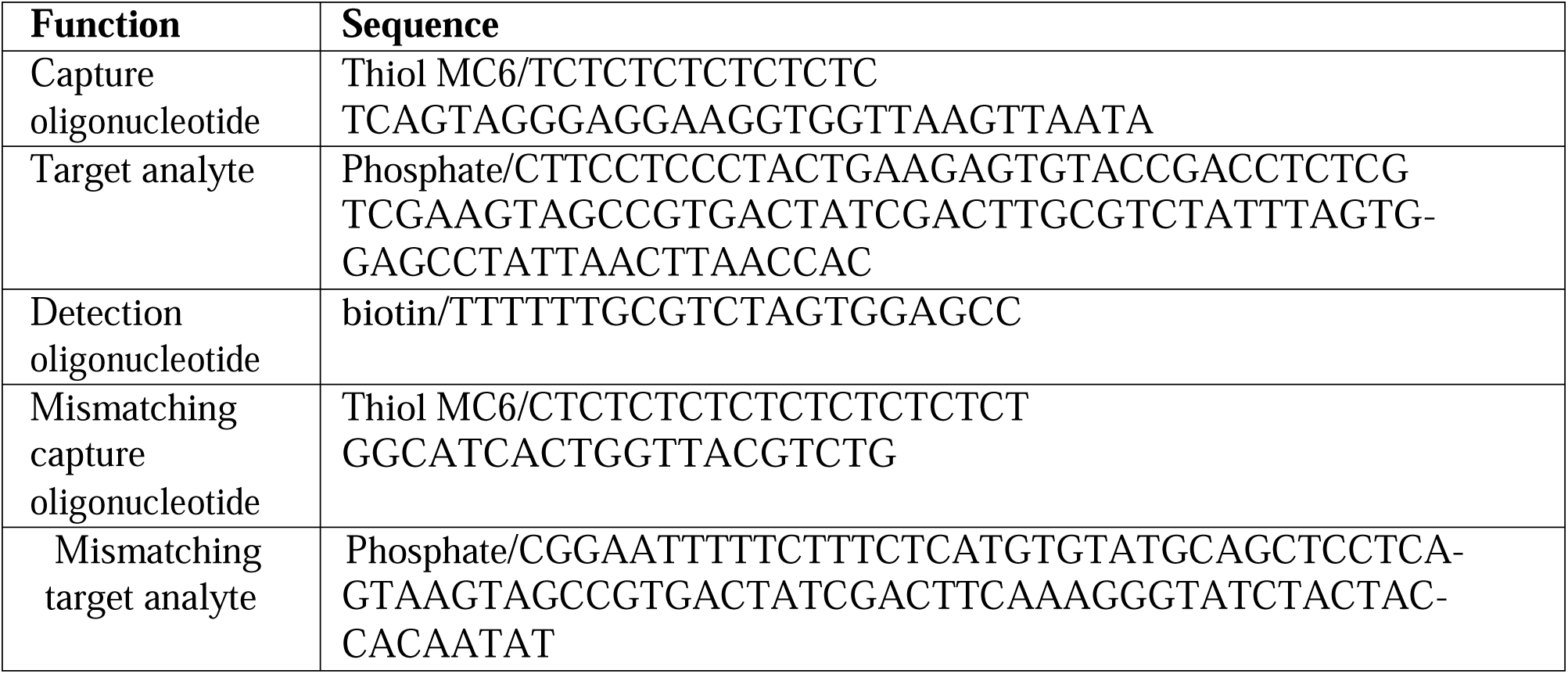
Oligonucleotide sequences used, written in 5’ → 3’ orientation

### Preparation of assay solutions

To prepare for the gold surface functionalization, 200 μL of a 20 μM solution of capture oligonucleotide solution was activated by incubation at RT for 1 h in freshly prepared buffer (pH 8.0) of 500 mM DTT and 10mM phosphate. After incubation, the solution was passed through a NAP-5 column to remove the DTT. The column was washed 5 times with 2 mL of Milli-Q water. Thereafter, 200 μL of the oligonucleotide solution was passed through the column, and 300 μL of Milli-Q water was added. The elution of the capture oligonucleotide solution was conducted by adding 500 μL of Milli-Q water. The surface functionalization solution was prepared as a mixture of 100 μM PEG, 10 % v/v SDS, 5 M NaCl, 100 mM phosphate buffer (pH 8), and 20 μM capture oligonucleotide solution in Milli-Q water.

Samples of varying concentration were prepared as follows. First we prepared a stock solution of DNA in Milli-Q water according to the DNA provider instructions for 100 μM concentrations. Thereafter we prepared diluted DNA in Milli-Q water solutions by serial 10× dilution. We mixed 5 μL of the diluted DNA solution with 5 μL of 10× phi29 polymerase buffer, 5 μL of 10 mM ATP, 1 μL of 20 μg/μL BSA, 0.5 μL of 5 U/μL T4 DNA ligase, and 5 μL of phi29 polymerase buffer, and added Milli-Q water until a total volume of 50 μL. The fraction of active substrate in the dilution series was determined to be 7.9 % (see supplementary information).

RCA amplification mix was prepared as a solution of 0.2 μg/μL BSA, 1X phi29 polymerase buffer, 250 μM dNTP, 200 mU/μL phi29 polymerase and 5 % v/v DMSO in Milli-Q water.

0.1 % DMSO buffer was prepared by a serial 10× dilution in Milli-Q water.

TNT rinsing buffer was prepared as a mixture of 10 mM of Tris-HCl and 150 mM of NaCl in Milli-Q water.

RCA rinsing buffer was prepared as a mixture of 10 % v/v 1× TNT and 1 % v/v of 10 % SDS in Milli-Q water.

We prepared hybridization buffer as a mixture of 1 M Tris-HCl pH 7.5, 0.5 M EDTA, 5 M NaCl, and 10 % Tween 20 in Milli-Q water. Streptavidin gold nanoparticles were functionalized with the detection oligonucleotide by mixing 17.7 % v/v streptavidin-functionalised gold nanoparticles solution, 50 nM detection oligonucleotide, 50 % v/v of 1 × hybridization buffer in Milli-Q water.

Gold enhancement buffer was prepared freshly from its constituents immediately before use according to the manufacturer instructions.

### Detector fabrication

Detectors consist of three layers: a top and bottom layer in OSTE+, which contain the wells, sandwiching the porous membrane (Fig. 4). First, a 5 nm/100 nm Ti/Au layer was deposited on both sides of the porous membranes by evaporation through a shadow mask under a 45° angle using a planetary mechanism, thus coating each well area and a separate top and bottom contact pad to each well. The evaporation angle avoids gold deposition on the entire walls of the pores. The top and bottom OSTE+ layers, both 1 mm thick, were structured using the OSTE+RIM process.^40^ After injection in an aluminum mold, the precursor was cured a first time by subsequent UV exposure using collimated light (300 mJ/cm^2^ at 365 nm) from a LS 30/7 1000 W NUV-light source (OAI, San Jose, CA, USA) for 60 s, waiting for 10 s, and curing for 20 s. The waiting prevents overheating of the polymer, thus preventing triggering of the second cure, which allows easy demolding of the layers from the mold after UV photocuring. After demolding, the two OSTE+ layers were aligned with the porous membranes and bonded during a second, thermal, cure at 90 ^°^C for 1 h. To functionalize the membranes, 300 μL of the capture oligonucleotide solution was added on each sample area and incubated overnight at 4 ^°^C.

**Figure 4:**
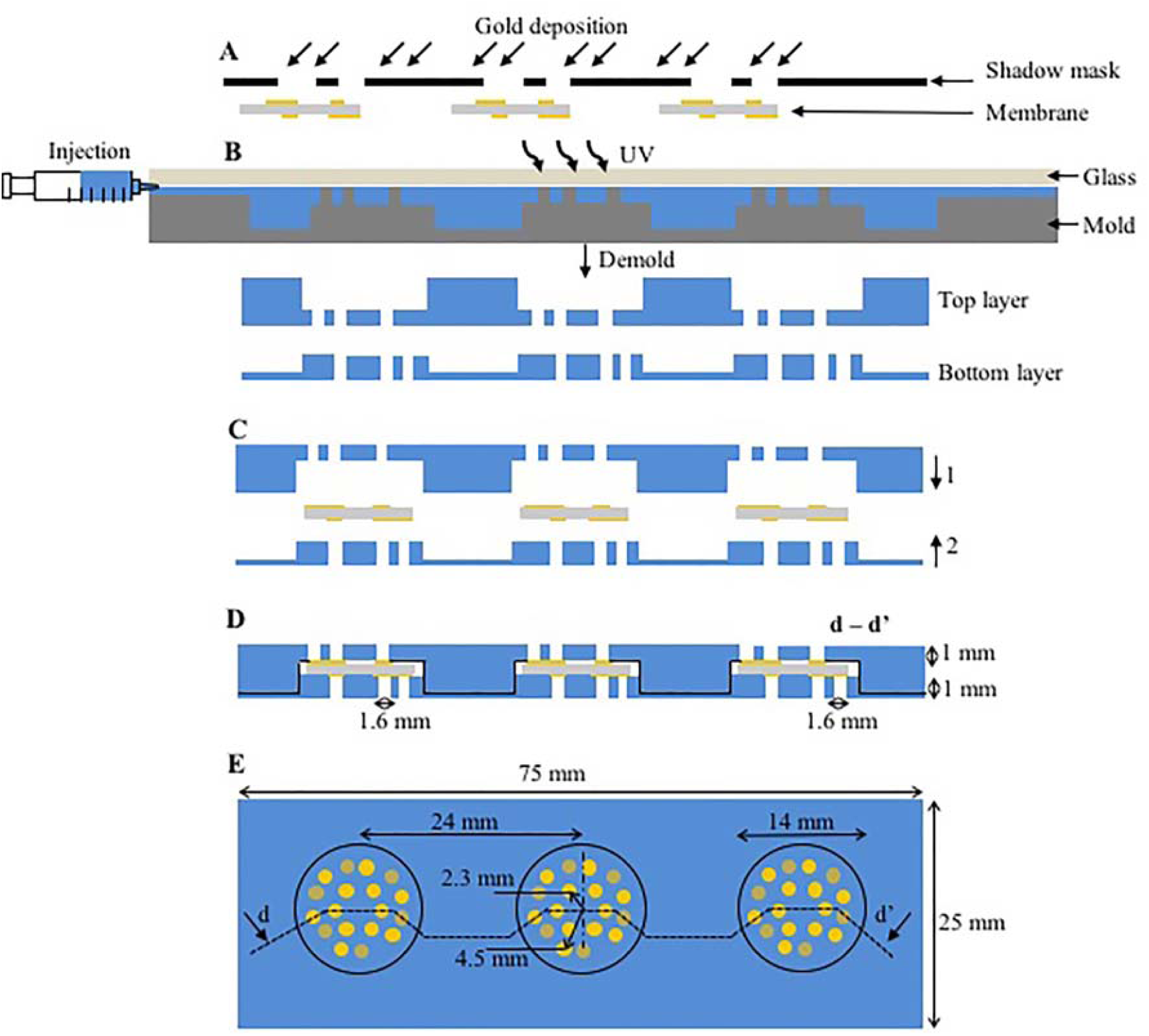
Fabrication and technical specification of the detector. A) A porous membrane is gold coated through a shadow mask. B) RIM of the OSTE+ top and bottom layer. C) Alignment and bonding of the three membranes between the top and bottom layer. D) Cross-sectional schematic of the detector. E) Top view schematic of the detector. OSTE+, off-stoichiometric thiol-ene-epoxy; RIM, reaction injection molding.

Two custom PDMS lids were built, each with three wells, that can be placed on the top and at the bottom of the detector to prevent evaporation during the RCA and gold nanoparticle incubation steps. First, an aluminum mold was milled (Minitech Machinery Corporation, Georgia, USA), then, both PDMS lids were casted in the mold and left to cure for 30 min at 80 ^°^C (Figure S4).

A custom vacuum chuck was built to provide suction at the bottom of every sample area during rinsing and DNA stretching steps. The vacuum chuck consisted of a holder with three holes, one for each sample area. A O-ring is placed surrounding each hole to avoid the issue of cross-contamination during each washing step. The holder was milled in PMMA (Figure S5).

### Detector operation

The detector operation was adapted from previous work.^20^

#### 1. Sample incubation (Fig. 1C)

Sample was added on top of each sample area in the following manner. First, 1 μL buffer was pipetted in the top and the bottom of every well to ensure no gas bubbles could block the flow through the wells. Second, a 50 μL sample droplet was added on the top of each sample area, covering all wells. The device was placed inside the PDMS lid (Figure S4) in a humid climate chamber at 37 ^°^C. The sample was left to flow through the membranes in the wells, driven by gravity, for a duration of *t*_*trans*_ = 35 min. Thereafter, no sample was visible on the top surface.

#### 2. Rinsing

The detector was placed on the vacuum chuck. 100 μL of TNT rinsing buffer was pipetted on the top of the sample areas. After all buffer had passed through all membranes, 100 μL of RCA rinsing buffer was pipetted on the top of each sample areas.

#### 3. RCA (Fig. 1D)

A 80 μL of RCA buffer droplet was added on the top surface of each sample area. The device was placed inside the PDMS lid (Figure S4) in a humid climate chamber at 37 ^°^C for 60 min.

#### 4. DNA stretching (Fig. 1E)

The detector was placed on the vacuum chuck. A total of 500 μL of 0.1 % DMSO buffer was pipetted on the top surface of the sample areas to cover the rest of RCA solution after RCA step. The vacuum chuck ensures a liquid-air interface travelling through the pores, from top to bottom, stretching the DNA concatemers.

#### 5. Rinsing

The detector was placed on the vacuum chuck. A total of 500 μL RCA rinsing buffer was pipetted on the top surface of each sample area in three additions.

#### 6. Gold nanoparticle incubation (Fig. 1F)

80 μL of gold nanoparticle buffer was added on the top surface of each sample area. The device was placed inside the PDMS lid in a humid climate chamber at 37 ^°^C for 60 min.

#### 7. Rinsing

The device was rinsed as described in step 5.

#### 8. Gold enhancement (Fig. 1G)

50 μL of gold enhancement solution was added on the top surface of each sample area. The device was placed inside the PDMS lid at RT for 60 min.

#### 9. Rinsing

Immediately after gold enhancement, the detector was placed on the vacuum chuck. 500 μL Milli-Q water was pipetted on the top surface of each sample area in three additions.

#### 10. Resistance measurement (Fig. 1H)

The trans-membrane resistance was measured for all wells using two-point probes (DY2000 Series Multi-Channel Potentiostat, Digi-Ivy Inc., Austin, TX, USA) contacting the contact areas. An electric potential was applied in the range 0–100 mV with a 2 mV step increase while measuring the current. The resistance was determined as the slope of the least-square linear fit to the I-V curve.

### SEM imaging

SEM images of the membrane top surfaces were acquired using a Hitachi SEM-Zeiss Ultra 55 instrument (ZEISS, Groupd headquarters, Oberkochen, Germany). An acceleration voltage of 5 kV was used during imaging of the membranes.

