## Supplemental Information for "Direct Electrical Detection of sub-aM DNA Concentrations"

### Supporting Information

#### **Determining the concentration of DNA substrate**

To determine the detector response curve, we are interested in the concentration of active substrate in the sample, i.e., we want to determine the actual concentration of RCA substrates by an independent measurement.

We performed RCA on the sample dilution series with varying oligonucleotide concentrations. Ligation and amplification were performed with the same reagent concentrations, and incubation times and temperatures as described in the methods section, with the exception of capture oligonucleotide (1 nM) and target analyte in varying concentrations (10 fM to 100 pM based on the initial concentration provided by the vendor, and confirmed by in-house Qubit fluorometer measurements). The assay was performed in solution in a tube by first ligating in 20 µL volume, adding 10 µL of amplification mixture and subsequently, fluorescently labelling the RCPs by adding 30 µL of hybridization buffer (1.4 M NaCl, 0.01% TWEEN 20, 20 mM Tris-HCl, pH 8, and EDTA) containing 5 nM Cy3-functionalized detection oligonucleotide DO for 2 min at 75 °C and 15 min at 55 °C.

5 µL of the RCA product (RCP) solutions were applied to, and uniformly spread on, a microscope slide over the coverslip area of 24 x 24 mm = 576 mm2. For imaging, a 20x magnification objective was used with a field of view of 0.665 x 0.665 mm = 0.44 mm2, corresponding to a volume of (0.44/576)x5 µL = 3.8 nL. For each concentration, images of three randomly chosen areas were acquired and the number of RCPs quantified using the open-source CellProfiler software (www.cellprofiler.org). Figure S3 shows the mean and of the resulting RCP count for each concentration. The measurement point for the 1 pM oligonucleotide concentration sample lays in the dynamic range, and indicates a mean of 180 RCPs $cps \sim\frac{180}{N_{A}\cdot3.8 nL}$ = 79 fM =7.9% of the designed concentration.

#### **Limitations of SEM imaging**

Whereas SEM imaging allowed qualitatively visualising the presence of gold nanowires on the membrane surface, we were neither able to quantify the nanowire density, nor to image trans-membrane gold nanowires in membrane cross-sections.

Quantitation of nanowire density on the membrane surface:

We observed a strong difference in number of wires between high and low concentration measurements. Unfortunately, wire counting on the sensor surface was practically not feasible. Every well contains a 2 mm^2^ membrane surface. To quantify wire formation at low concentrations, these large membranes would need to be entirely scanned in a SEM at very high magnification. Already on a flat membrane surface, this would be a very tedious and time-consuming effort. In our sensor, the membranes are suspended deep inside mm-scaled pores. The sidewalls of the pores give rise to substantial charging effects in the SEM (even after substantial gold sputtering of the samples), making high magnification views very difficult, i.e., the image apparently continuously “shifts” during SEM rastering. The capture of a single SEM image with a field of view of a few tens of micrometre, e.g. as depicted in Fig. 1H, easily requires 30 min of trial and error. It is thus practically infeasible to scan even a small portion of the membrane for wire counting.

Visualisation of trans-membrane gold nanowires in membrane cross-sections:

Three strategies (i-iii) were attempted to visualise the nanowires in cross-sectional SEM studies.

i) Cutting the membrane using a normal scissor resulted in gold layer removal and was thus not feasible.

ii) Cutting the membrane using very low power laser also resulted in gold layer removal and additionally in burning of the membrane despite the low laser power.

iii) Freezing the membrane at -80 C followed by microtome cutting and 10 nm gold deposition allowed visualising the membrane cross-sections. Figure S7 shows typical results.

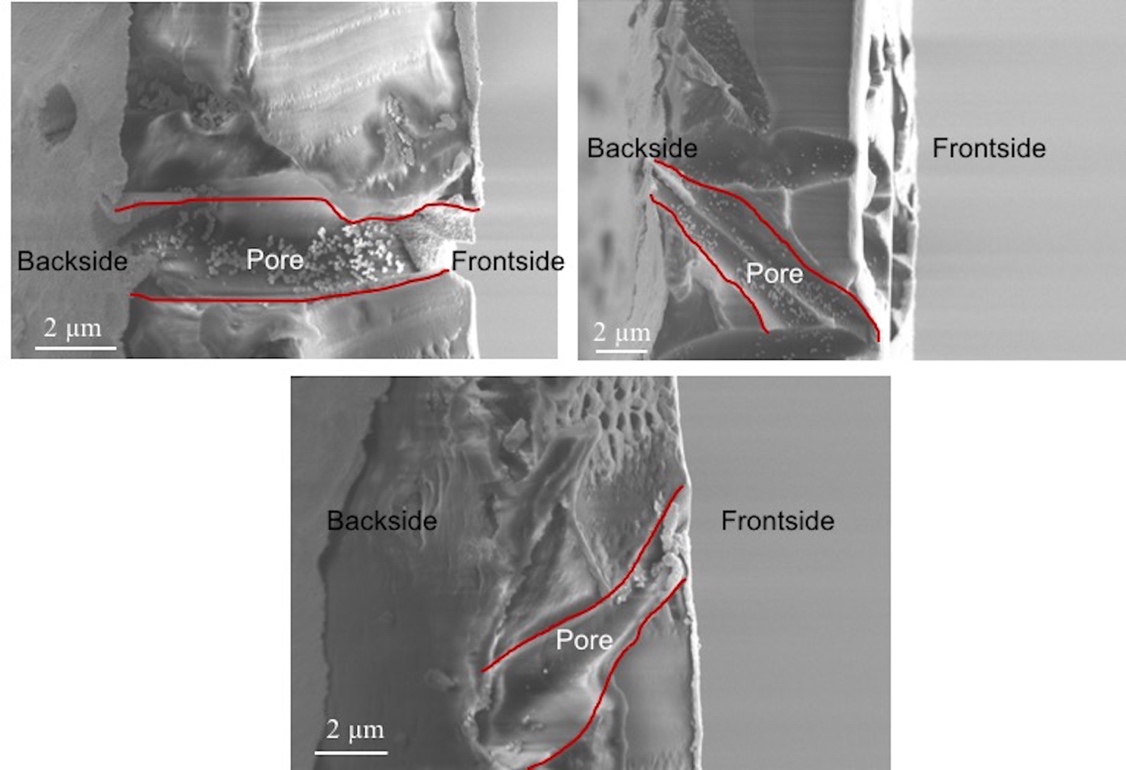

Figure S7. Three side images of membranes cut after freezing a -80 C. The micropores were heavily distorted and we could not find back features that looked like nanowires.

We found substantial distortion of the membrane and its pores. However, we could not observe any gold wires, neither on the membrane surface in proximity of the cut, nor in the pores. We attribute this to the breaking off of the gold wires during cutting and material distortion.

Focused ion beam milling (FIB) was considered, but discarded given the expected burning of the membrane and the re-deposition of debris during this process, which would make interpretation of images likely impossible.

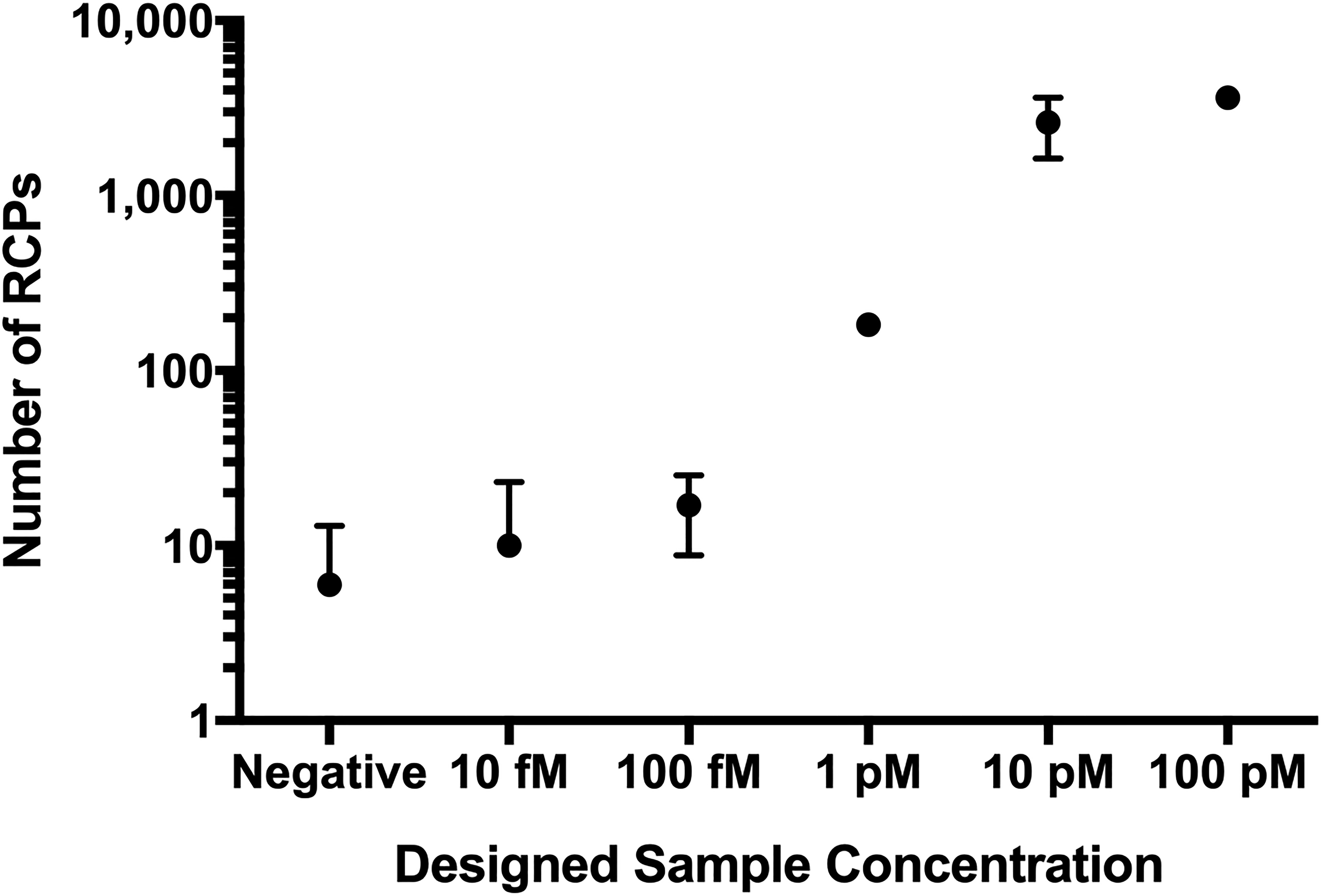

Figure S3: Counted number of RCPs versus the designed sample concentration.

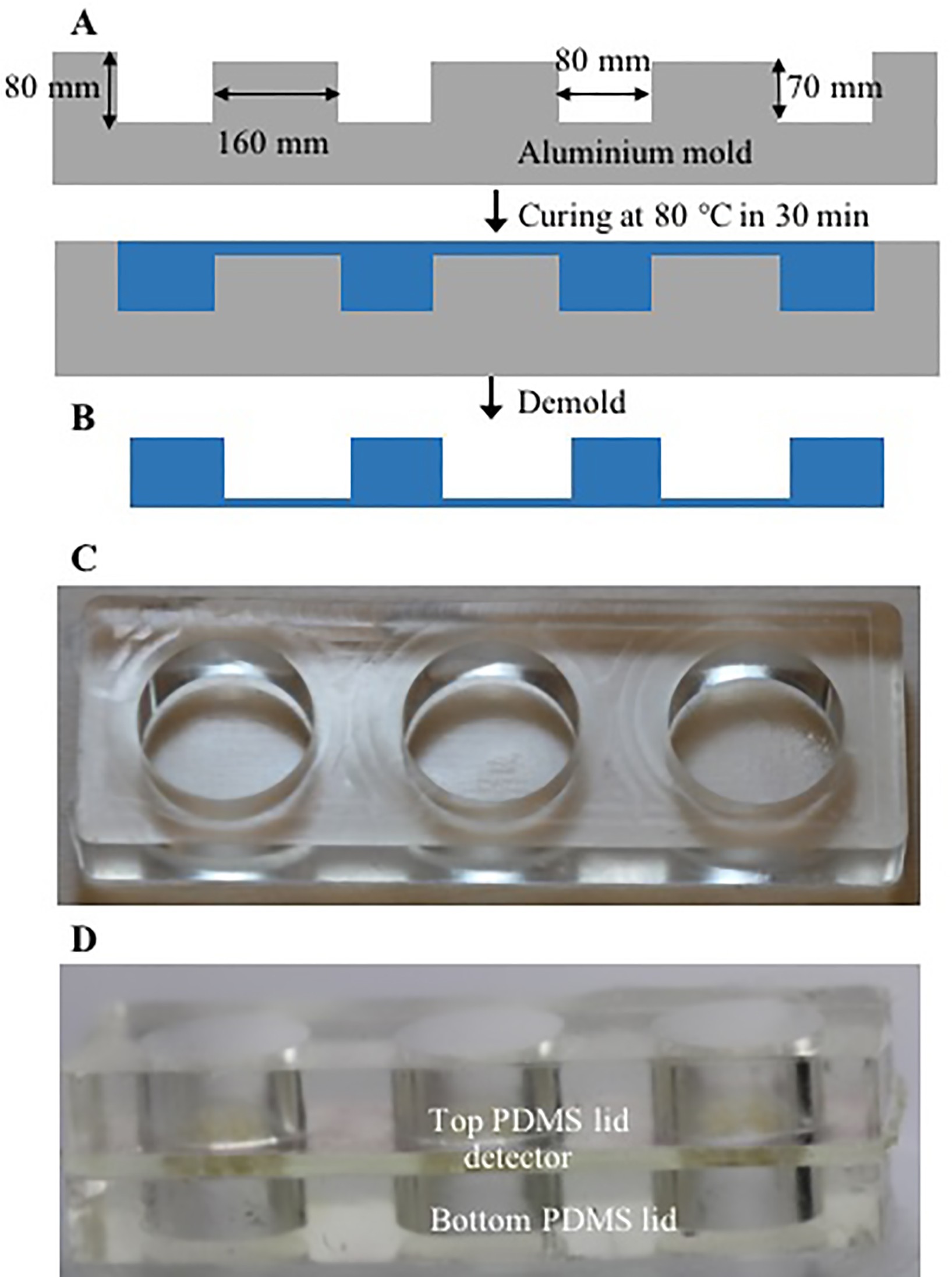

Figure S4: A) Cross-sectional schematic of the aluminum mold, and B) the PDMS casting process, C) photograph of the resulting PDMS lid, and D) photograph of a detector between a top and bottom lid for prevention of evaporation during RCA and gold nanoparticle enhancement.

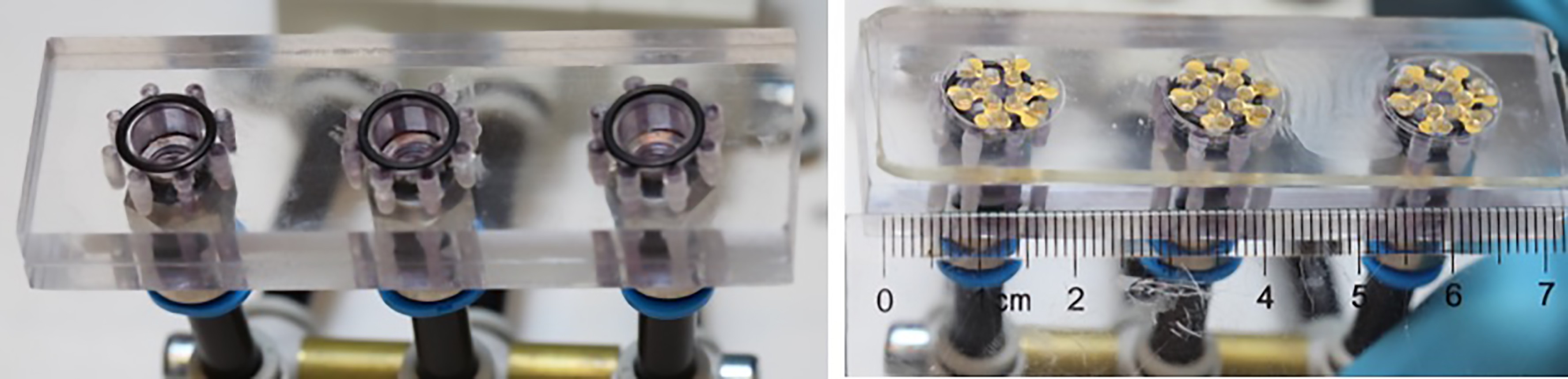

Figure S5: Custom vacuum chuck to provide fluid actuation through the detector membranes. Chuck without detector (left) and with detector on top (right).

Table S3: List of the resulting trans-membrane resistances for each well of eight detector measurements vs the active sample concentration. Open circuit results are denoted. The centre column shows the result for varying sample concentrations. The left column shows the positive control results (790 fM). The right column shows the negative control results (79 or 7.9 zM).

∞

| 790 fM | 7.9 aM | 7.9 zM |
| --- | --- | --- |
| 40.9 Ω | 28.8 Ω | ∞  ∞  ∞  ∞  ∞  ∞ |
| 24.2 Ω | 18.9 Ω |  |
| 21.7 Ω | 32.6 Ω |  |
| 23.6 Ω | 107.1 Ω |  |
| 31.2 Ω | 32.5 Ω |  |
| 27.9 Ω | ∞ |  |
| 790 fM | 7.9 aM | 7.9 zM |
| 25.3 Ω | 625.0 Ω | ∞  ∞  ∞  ∞  ∞  ∞ |
| 25.0 Ω | 28.8 Ω |  |
| 20.7 Ω | 18.8 Ω |  |
| 18.3 Ω | 89.4 Ω |  |
| 25.8 Ω | 24.4 Ω |  |
| 106.3 Ω | 26.0 Ω |  |
| 790 fM | 790 zM | 79 zM |
| 25.0 Ω | 58.3 Ω | ∞  ∞  ∞  ∞  ∞  ∞ |
| 20.5 Ω | 21.1 Ω |  |
| 18.4 Ω | 35.6 Ω |  |
| 25.0 Ω | 58.3 Ω |  |
| 106.6 Ω | 46.7 Ω |  |
| 27.6 Ω | 37.5 Ω |  |
| 790 fM | 790 zM | 79 zM |
| 20.0 Ω | 666.7 Ω | ∞  ∞  ∞  ∞  ∞  ∞ |
| 18.2 Ω | 61.0 Ω |  |
| 86.1 Ω | 274.7 Ω |  |
| 15.9 Ω | 21.5 Ω |  |
| 22.1 Ω  14.1 Ω | ∞  ∞ |  |

| 790 fM | 7.9 aM | 7.9 zM |
| --- | --- | --- |
| 41.5 Ω | 69.1 Ω | ∞  ∞  ∞  ∞  ∞  ∞ |
| 39.4 Ω | 21.7 Ω |  |
| 40.0 Ω | 102.1 Ω |  |
| 36.7 Ω | 749.6 Ω |  |
| 31.3 Ω | 666.7 Ω |  |
| 25 Ω | 294.1 Ω |  |
| 790 fM | 7.9 aM | 7.9 zM |
| 28.0 Ω | 16.1 Ω | ∞  ∞  ∞  ∞  ∞  ∞ |
| 36.2 Ω | 56.2 Ω |  |
| 129.2 Ω | 23.8 Ω |  |
| 27.7 Ω | 26.0 Ω |  |
| 129.3 Ω | 37.6 Ω |  |
| 13.2 Ω | ∞ |  |
| 790 fM | 790 zM | 79 zM |
| 64.1 Ω | 249.1 Ω | ∞  ∞  ∞  ∞  ∞  ∞ |
| 32.4 Ω | 208.8 Ω |  |
| 18.8 Ω | 86.5 Ω |  |
| 45.2 Ω | 35.4 Ω |  |
| 13.2 Ω  32.5 Ω | ∞  ∞ |  |
| 790 fM | 790 zM | 79 zM |
| 36.0 Ω | 20.1Ω | ∞  ∞  ∞  ∞  ∞  ∞ |
| 129.3 Ω | 31.4Ω |  |
| 27.5 Ω | 56.8 Ω |  |
| 25.1 Ω | 46.7 Ω |  |
| 107.2 Ω | 22.6 Ω |  |
| 98.5 Ω | ∞ |  |

Table S4: Detector response for different control experiments. Control A - Mismatched target (padlock probe); Control B - Mismatched DNA receptor; Control C - Without DNA Receptor; and Control D - Omitting AuNPs. Positive control, mismatching target and receptor were used at a concentration of 10 pM. The high resistance/open circuit values of the controls demonstrate the high specificity of the detector.

| **Positive Control** | **Control A** | **Control B** | **Control C** | **Control D** |
| --- | --- | --- | --- | --- |
| 39.0 Ω | *>* 108 Ω  ∞  ∞  ∞  ∞  ∞ | ∞  ∞  ∞  ∞  ∞  ∞ | ∞  ∞  ∞  ∞  ∞  ∞ | *>* 108 Ω  ∞  ∞  ∞  ∞  ∞ |
| 15.1 Ω |  |  |  |  |
| 54.5 Ω |  |  |  |  |
| 39.1 Ω |  |  |  |  |
| 22.0 Ω |  |  |  |  |
| 37.6 Ω |  |  |  |  |

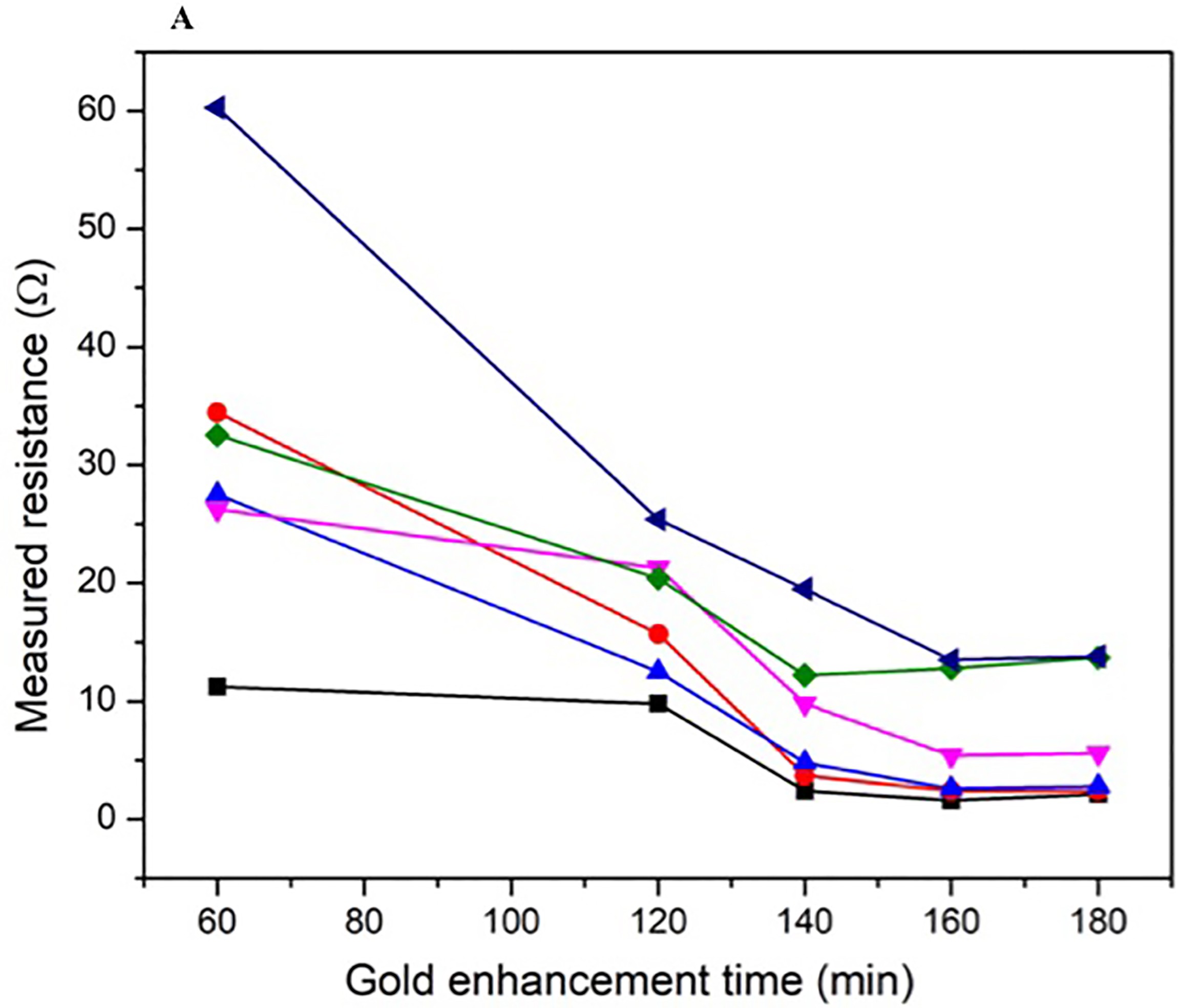

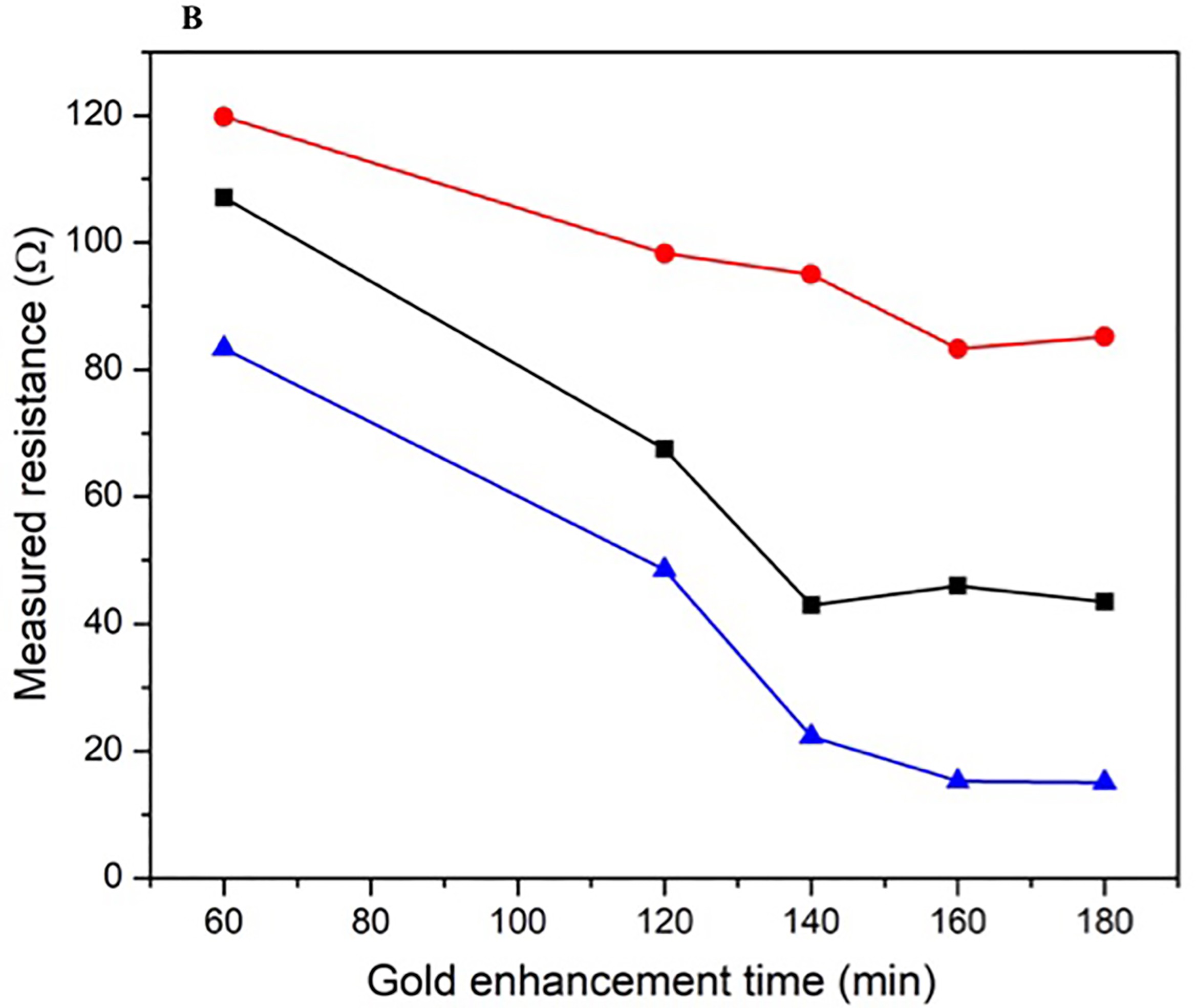

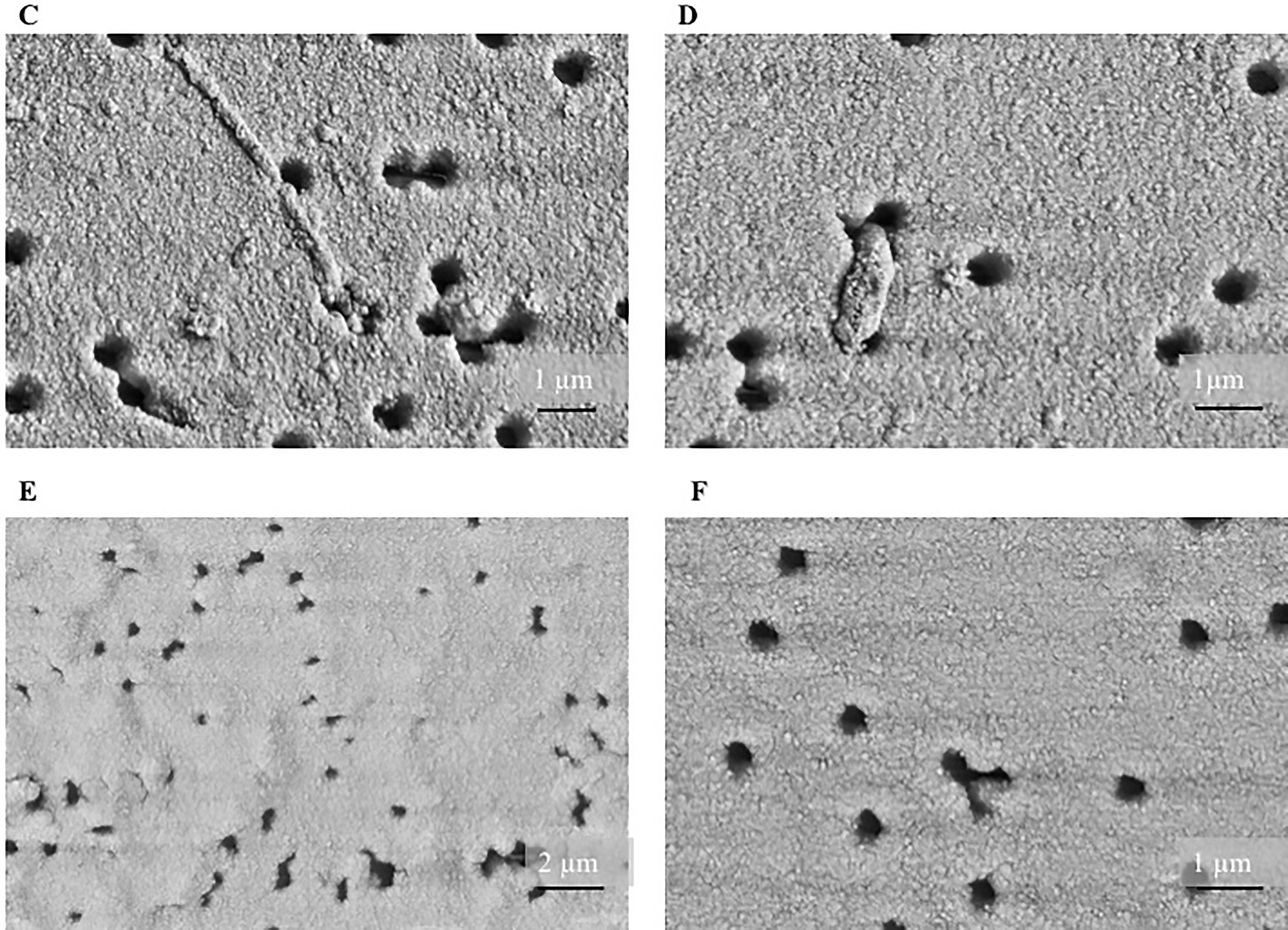

Figure S6: The trans-well resistance at different gold enhancement times for A) 790 fM and B) 790 zM sample concentrations. Lines connect measurement points for one well and are added for eye guidance. After 160 min enhancement, the resistance value does not change significantly. SEM image of the top surface for C) 790 fM and D) 790 zM sample concentrations, and of the bottom surface for E) 790 fM and F) 790 zM sample concentrations after 180 min enhancement. The pore diameters are reduced to 500 nm and 200 nm, respectively. The resulting fluidic resistance (~1*/R_p_*^4^) has thus increased with an estimated 16-fold to 625-fold. One can expect a similar reduction in gold enhancement solution transport, which explains the saturation of the resistance values.
